# A photoreceptor-based hydrogel with red light-responsive reversible sol-gel transition as transient cellular matrix

**DOI:** 10.1101/2023.04.04.535523

**Authors:** Maximilian Hörner, Jan Becker, Rebecca Bohnert, Miguel Baños, Carolina Jerez-Longres, Vanessa Mühlhäuser, Daniel Härrer, Tin Wang Wong, Matthias Meier, Wilfried Weber

## Abstract

Hydrogels with adjustable mechanical properties have been engineered as matrices for mammalian cells and allow the dynamic, mechano-responsive manipulation of cell fate and function. Recent research yielded hydrogels, where biological photoreceptors translated optical signals into a reversible and adjustable change in hydrogel mechanics. While their initial application provided important insights into mechanobiology, broader implementation is limited by a small dynamic range of addressable stiffness. Here, we overcome this limitation by developing a photoreceptor-based hydrogel with reversibly adjustable stiffness from 800 Pa to the sol state. The hydrogel is based on star-shaped polyethylene glycol, functionalized with the red/far-red light photoreceptor phytochrome B (PhyB), or phytochrome-interacting factor 6 (PIF6). Upon illumination with red light, PhyB heterodimerizes with PIF6, thus crosslinking the polymers and resulting in gelation. However, upon illumination with far-red light, the proteins dissociate and trigger a complete gel-to-sol transition. We comprehensively characterize the hydrogel’s light-responsive mechanical properties and apply it as reversible extracellular matrix for the spatiotemporally controlled deposition of mammalian cells within a microfluidic chip. We anticipate that this technology will open new avenues for the site- and time-specific positioning of cells and will contribute to overcome spatial restrictions.

## 1. Introduction

Traditionally, *in vitro* cell culture is performed on two-dimensional substrates like polystyrene, and in the past, studies under these artificially defined conditions provided the basis for our fundamental understanding of cellular processes like differentiation, proliferation, and migration. However, it was demonstrated that cellular fate and function are also dependent on the mechanical properties of the culture substrate.^[1]^ *In vivo*, cells are often embedded in soft tissues that exhibit significantly lower mechanical stiffness than conventional *in vitro* systems^[2]^ and their interactions with this highly dynamic 3D extracellular matrix (ECM) are crucial for tissue development^[3]^ and subsequent homeostatic control of its structure and function.^[4]^ Monolayer cultivation and removal of cells from this natural microenvironment can result in aberrant behavior^[5]^, which may lead to misleading or incomplete conclusions. Therefore, novel materials with reversibly and dynamically adjustable mechanical stiffness are urgently demanded to overcome the current limitations and provide a robust platform for further advances in basic research and medical applications. A promising model among these materials are hydrogels, 3D crosslinked polymer networks, that can absorb and retain a large amount of water, making them highly biocompatible and ideal as a scaffold for cell growth, delivery of drugs and other biological molecules, as well as tissue engineering.^[6,7]^

Recent advances in material sciences and synthetic biology generated a variety of smart hydrogel designs that allow for dynamic modulation of selective mechanical properties and therefore more closely resemble the *in vivo* environment.^[8–12]^ For example, materials responsive to external stimuli of chemical nature (pH^[13]^ or active molecules^[14]^) or physical cues (temperature^[15]^, light^[16,17]^, or magnetic fields^[18]^) were developed. Hydrogels that allow modulation of their mechanics over time can present a versatile opportunity to establish novel dynamic cell culture systems.^[17,19–21]^ By further combining such stimuli-responsive hydrogels with microfluidic systems, the experimental microenvironment can be defined in greater detail and novel technologies, that maximize their individual advantages, can emerge.^[22]^ For the successful integration of hydrogels into microfluidics, and to benefit from the conveniences of ‘lab-on-a-chip’ experiments, high spatiotemporal precision is required, since microfluidics are per definition applied in the micro-to nanometer scale. Desired characteristics for an ideal hydrogel, therefore, include the opportunity to control its properties at high spatiotemporal resolution, using non-invasive stimuli to address a wide dynamic range with intermediate states via a reversible and robust mechanism.

The optimal stimulus for inducing such alterations is light, as it can be applied in a reversible, adjustable, and local manner at excellent resolution.^[23]^ In order to translate optical input into changes in the hydrogels’ mechanical properties, chemical or biological photo-responsive moieties are required. For such applications, biological photoreceptors show highly suitable features as they are inherently functional in a biological background, are responsive to the biocompatible light spectrum and intensities, and show high specificity, even in a complex biochemical environment. Of special interest for modulating hydrogel properties are photoreceptors that dimerize in the presence of light, either by forming homodimers or heterodimers with another binding partner. The coupling of such photoreceptors to polymers allows light-responsive polymer crosslinking and thus stiffening of the scaffold. Upon dissociation of the photoreceptors, polymer crosslink density decreases and the material softens. Several photoreceptor-based hydrogels have been constructed and yielded important insights into mechano-signaling or were used to control migration of cells (for reviews, see ^[12,16]^).

Despite these impressive advances, the broader application of these hydrogels is limited due to the requirement for energy-rich light and/or a low dynamic range. For example, light-responsive hydrogels based on the photoreceptors UVR8^[24,25]^ or LOV2^[26]^ require UV or blue light and therefore exhibit low tissue penetration^[27]^ and possible cytotoxic effects.^[28,29]^ Other materials suffer from a relatively low dynamic range (EL222^[30]^), limited reversion cycles (Dronpa145N^[31,32]^), or other drawbacks like instability or irreversibility (CarHC^[33,34]^ and PhoCl^[35]^). For a comparison of these hydrogels with regard to their dynamic range and illumination requirements, see a recent review article.^[12]^ In the past, we published a revised design of a light-controlled hydrogel based on the cyanobacterial phytochrome receptor Cph1 conjugated to branched polyethylene glycol (PEG).^[36–39]^ However, these gels were limited to a 2-3-fold change in material stiffness.

In this study, we introduce an enhanced photoreceptor-based hydrogel design that overcomes the above-described limitations. The incorporated photoreceptor is part of the phytochrome (Phy) class which are capable of absorbing light in the red/far-red spectrum, making them more suitable for biological application due to increased tissue penetration and lower spectral energy of red/far-red light.^[40–42]^ The scaffolding polymer PEG, has been clinically approved in combinations with pharmaceuticals to increase their circulation half-life and is a commonly used biocompatible and non-immunogenic synthetic polymer in hydrogels.^[43,44]^ In this design, star-shaped PEG is reversibly crosslinked by heterodimerization of *Arabidopsis thaliana* derived phytochrome B (PhyB) with the phytochrome interacting factor 6 (PIF6). Red light of 660 nm wavelength leads to a cis-trans isomerization of the covalently bound chromophore phycocyanobilin (PCB) and transfers PhyB into its active P_fr_ state, which selectively binds PIF6.^[45]^ Illumination with far-red light of 740 nm reverts the phytochrome into its inactive, red light absorbing P_r_ state. By coupling PhyB or PIF6 to 4-arm or 8-arm PEG-vinyl sulfone (PEG-VS), we synthesized a heterodimeric polymer system that is crosslinked under 660 nm and dissociated under 740 nm illumination. These alterations in interaction trigger a fully reversible and adjustable change in the storage modulus from approximately 800 Pa down to the sol state. We further applied these light-reversible sol-gel transitions to optically pattern cells within a microfluidic chip.

## 2. Results and discussion

### 2.1 Design of the hydrogel with photo-reversible gel-sol transition

In order to generate switchable hydrogels based on photo-reversible polymer crosslinking, we decorated star-shaped PEG polymers either with PhyB via Spy chemistry, or with PIF6 by Michael-type addition (Figure 1). Spy chemistry describes a system consisting of the small peptide SpyTag (13 amino acids (AAs)) and the protein SpyCatcher (116 AAs) that spontaneously and rapidly form an isopeptide bond to covalently couple two proteins of interest.^[46]^

**Figure 1:**
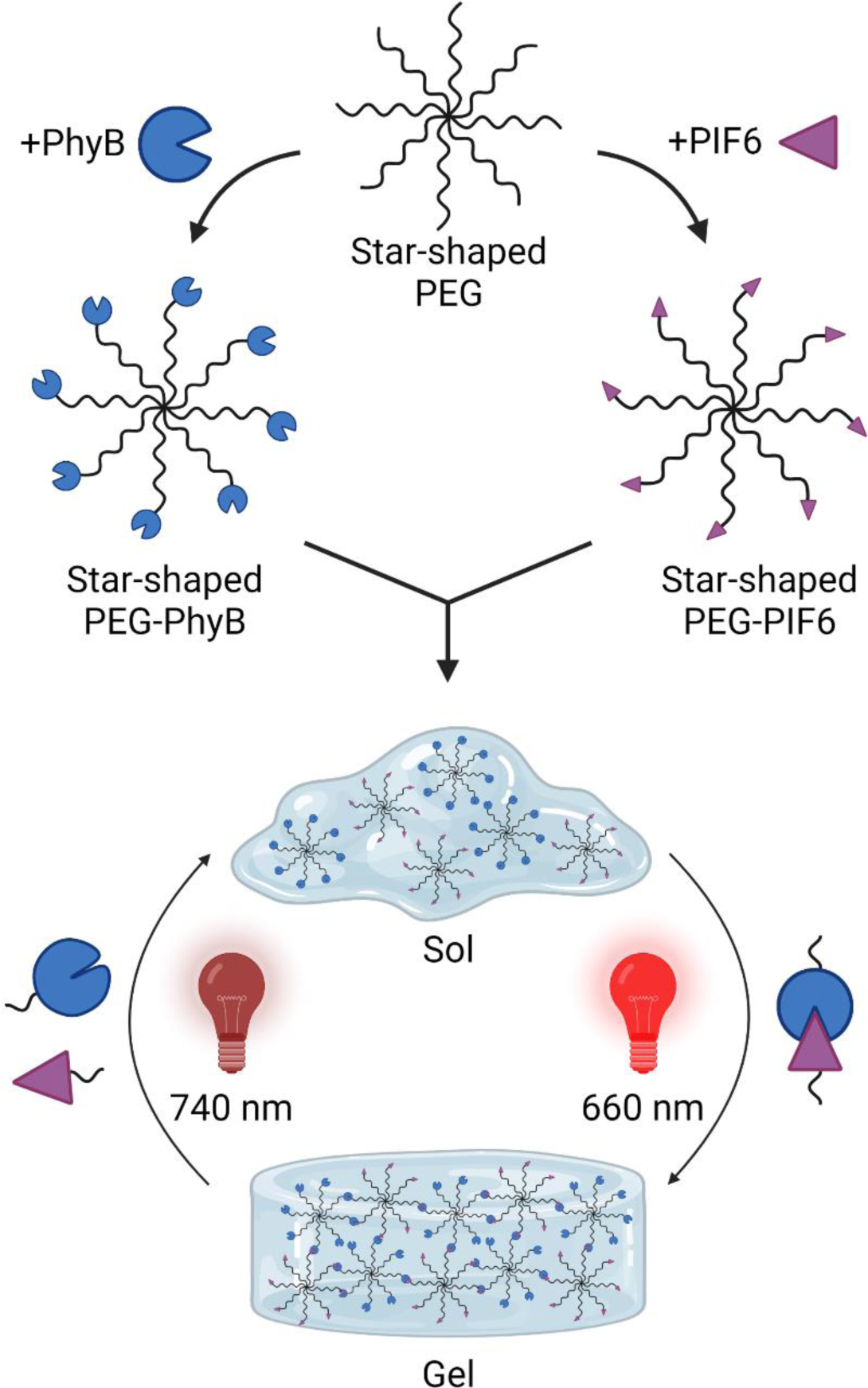
Hydrogel design and functionality. Star-shaped polyethylene glycol (PEG) is decorated either with *Arabidopsis*-derived phytochrome B (PhyB) or phytochrome interacting factor 6 (PIF6). Red light application of 660 nm wavelength induces heterodimerization of the two polymers, causing polymerization and hydrogel formation. This process is reversed by illumination with far-red light of 740 nm to transform the hydrogel back into its original sol state.

As a result, two types of protein-coupled polymers, that exhibit a light dependent interaction mechanism, were synthesized. When combined, 660 nm light induced a conformational change within PhyB and transferred it into its active P_fr_ state, which selectively binds PIF6 in a non-covalent manner. Hence, the crosslinking density between PEG-PhyB and PEG-PIF6 was drastically enhanced, leading to an increase in material stiffness and the formation of a hydrogel. This process could be fully reverted by application of far-red light (740 nm) to transform PhyB into its inactive non-binding P_r_ state.

### 2.2 Synthesis and characterization of hydrogel precursors

Since Spy chemistry offers a highly selective and gentle coupling process, we implemented this system for PhyB conjugation to PEG. The individual steps of this process are schematically illustrated in **Figure 2A**. First, a SpyTag peptide with a cysteine (Cys) residue was synthesized and coupled to PEG-VS (40 kDa, 8-arm) via Michael-type addition, resulting in the formation of a thioether bond. Uncoupled peptide was subsequently removed by dialysis, as confirmed by the subsequent size exclusion chromatography (SEC) analysis at 280 nm absorbance (Figure 2B), resulting in a peptide-coupled polymer.

**Figure 2:**
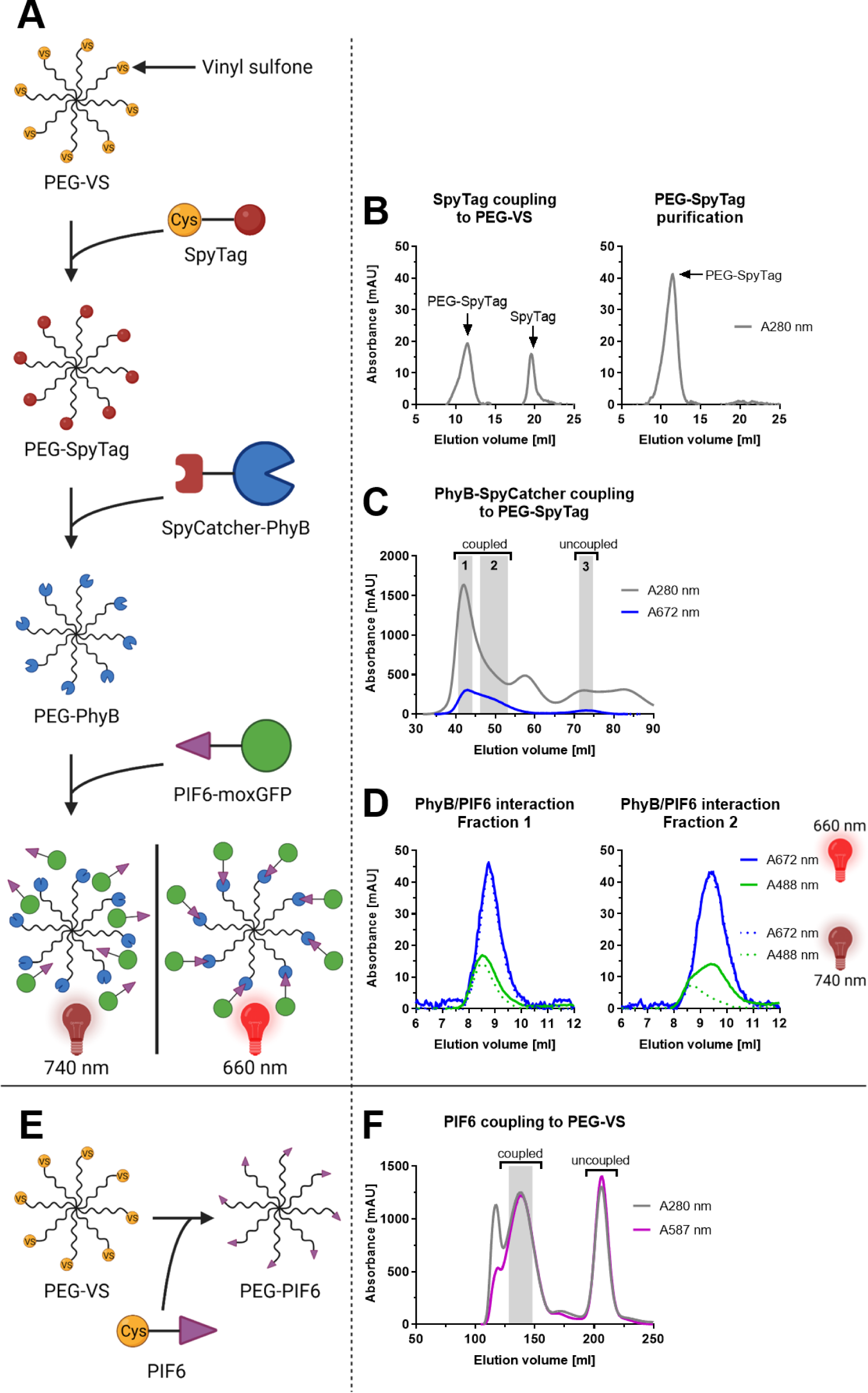
Coupling of PhyB and PIF6 to PEG-VS, as well as functional characterization of PEG-PhyB. (A) Schematic illustration of PhyB conjugation to PEG-VS via Spy chemistry, as well as light-controlled interaction of PEG-PhyB with moxGFP-PIF6. (B) SpyTag coupling to PEG-VS. SpyTag peptide and PEG-VS were incubated overnight and coupling efficiency was analyzed with size exclusion chromatography (SEC). Afterwards, the product was dialyzed against water to remove uncoupled SpyTag and purity was assessed by SEC. (C) PhyB-SpyCatcher (PhyB-Spy) coupling to PEG-SpyTag. Purified PhyB-Spy protein was incubated with PEG-SpyTag for 2 h before purification of the conjugate by SEC (detection at 280 nm (protein) and 672 nm (PhyB)). Fractions were collected and pooled as indicated by the grey areas; fraction 2 was used for later hydrogel synthesis. (D) Light-dependent PEG-PhyB interaction with moxGFP-PIF6. Fraction 1 or 2 of PEG-PhyB SEC (see C) were mixed with purified moxGFP-PIF6 protein and illuminated for 2 min with red (660 nm, solid lines) or far-red (740 nm, dotted lines) light before separation by SEC and analysis of 488 nm (moxGFP-PIF6) and 672 nm (PhyB) absorbance. (E) Schematics of PIF6 conjugation to PEG-VS via Michael-type addition. (F) PIF6 coupling to PEG-VS. PEG-VS was incubated with purified mCherry-PIF6 protein (N-terminal Cys residue, referred to as PIF6) overnight and the conjugate was purified by SEC (absorbance was measured at 280 nm (protein) and 587 nm (PIF6)). Fractions of the high molecular weight peak (indicated in grey) were pooled and used for later hydrogel synthesis.

Next, the fusion protein PhyB-SpyCatcher (PhyB-Spy) was engineered by cloning the SpyTag-binding domain SpyCatcher downstream of the photosensory domain of PhyB (AAs 1-651) from *A. thaliana*, followed by a His_6_-Tag for protein purification (Figure S1). The coding sequence of PhyB-Spy was placed in an expression vector that further harbors the genes *ho1* and *pcyA* for biosynthesis of the chromophore PCB. PhyB-Spy was recombinantly produced in *E. coli* and subsequently purified by immobilized metal affinity chromatography (IMAC). The sodium dodecyl sulfate-polyacrylamide gel electrophoresis (SDS-PAGE) of the purified protein revealed a distinct band with the expected size (calculated molecular weight of PhyB-Spy: 85 kDa) and the successful conjugation of PCB was verified by a Zn^2+^ staining of the gel (Figure S2A).^[47]^

Afterwards, we experimentally determined the optimal coupling ratio of PhyB-Spy and PEG-SpyTag. To this aim, PhyB-Spy and SpyTag-functionalized polymers were mixed at different protein:polymer ratios, ranging from 0.1 to 10, and the coupling efficiencies were analyzed by SDS-PAGE (Figure S3A). The optimal ratio is a trade-off between efficient decoration of polymers with PhyB-Spy while minimizing the loss of uncoupled protein and was determined to be PhyB-Spy:PEG-SpyTag = 2:1.

In the following, PhyB-Spy and PEG-SpyTag were coupled at larger scale with the previously determined optimal ratio and the coupling was analyzed by SEC with monitoring protein absorbance at 280 nm and PhyB specific absorption at 672 nm (isosbestic point of PhyB) (Figure 2C). The latter displayed a main peak with a high molecular weight which we hypothesized to be the coupled protein-polymer fraction, as well as a significantly weaker peak with lower molecular weight, presumably uncoupled PhyB-Spy protein. Although, the Spy coupling chemistry is highly specific, we found that the main peak could be further divided into two different fractions (indicated by grey bars in Figure 2C) of protein-polymer conjugates that differ from each other in the ratio of 280 nm: 672 nm absorbance. Both fractions of the main peak were further characterized and it was analyzed whether they contained proteins crosslinked by disulfide bridges. To this aim, a SDS-PAGE analysis under reducing and non-reducing conditions was performed which revealed that among the coupled fractions of the main peak, the fraction with the lower 280 nm: 672 nm ratio (fraction 1) contained large amounts of proteins covalently attached by disulfide bridges.

To assess the functionality of PhyB in the two PEG-PhyB fractions, we analyzed the light-dependent interaction with its interaction factor PIF6 by analytical SEC after illumination with 740 nm or 660 nm light. For this approach, we designed, produced, and purified the fusion protein moxGFP-PIF6 (Figure S1 and S2B). The absorption maximum of moxGFP at 488 nm is within the absorption minimum of PhyB, which allowed specific detection of both proteins during SEC by measuring the absorbance at 488 nm and 672 nm. We observed only for fraction 2 a light-dependent interaction between moxGFP-PIF6 and PEG-PhyB, whereas fraction 1 of PEG-PhyB interacted in a light-independent manner with moxGFP-PIF6 (Figure 2D). Therefore, only fraction 2 of the PEG-PhyB purification was used for later hydrogel synthesis. To couple PIF6 to 8-arm PEG, we engineered a fusion protein consisting of a Cys residue, the fluorescent protein mCherry, and PIF6 (AAs 1-100, C9S, C10S) from *A. thaliana* (Figure S1). We introduced the two cysteine-to-serine mutations in PIF6 to allow specific coupling to PEG-VS solely via the newly introduced Cys residue at its N-terminus. This fusion protein (referred to as PIF6) was produced in *E. coli* and purified by IMAC, resulting in a pure and monomeric protein of the expected size (calculated molecular weight: 40 kDa), as verified by SDS-PAGE and SEC (Figure S2C). We coupled PIF6 to PEG-VS via Michael-type addition of the Cys residue to the vinyl sulfone group of the branched PEG (Figure 2E) and purified the conjugate PEG-PIF6 by SEC (Figure 2F).

### 2.3 Hydrogel synthesis and characterization

After the development of the conjugation process of PhyB and PIF6 to star-shaped PEG and the subsequent purification strategy, we next evaluated the light-controlled hydrogel formation by mixing purified PEG-PhyB and PEG-PIF6. For this step, and the later application in the presence of mammalian cells, we further included the cell adhesion motif RGD at the C-terminus of PhyB and used a truncated SpyCatcher lacking short AA stretches at its N- and C-terminus that were previously shown to interact with different cell surface components in a non-specific manner.^[48]^ The resulting protein (PhyB-SpyΔ) was produced and characterized as before (Figure S1 and Figure S4). In addition, we investigated different formulations by evaluating the influence of two different star-shaped PEG molecules, 20 kDa 4-arm and 40 kDa 8-arm PEG on the hydrogels’ mechanical properties.

First 4-arm or 8-arm PEG-PhyB were mixed with 8-arm PEG-PIF6 (final coupled PhyB concentration: 50 mg/ml) under illumination of 740 nm. Illumination with 660 nm light resulted in gelation within seconds and the materials were subjected to rheological characterization via small amplitude oscillatory shear measurements (**Figure 3A**). In this method the storage (G’) and loss moduli (G’’) were recorded to provide information about the elastic and viscous part of the sample, respectively. In general, a sample is considered to exhibit gel properties as long as G’>G’’ holds true.^[49]^ The measurements revealed that the hydrogel with 4-arm PEG-PhyB was approximately 1.8-fold-stiffer with G’ values of 409 ± 4 Pa compared to 225 ± 6 Pa of the hydrogel with 8-arm PEG-PhyB. We hypothesize that in the 4-arm PEG-PhyB hydrogels more of the relatively large PhyB proteins can participate in the interaction with PIF6 due to reduced steric hindrance and thus increased flexibility, manifesting in a higher mechanical stiffness. After illumination with 740 nm light to investigate the light responsiveness, we observed that both hydrogel variants responded with an immediate decrease in stiffness that reached its plateau within a 20 s time frame. The recorded storage modulus dropped to the levels of the loss modulus (<10 Pa for both variants), indicating the dissolution of the hydrogel as demonstrated by the G’’:G’ ratio ≈ 1 (tan(δ)). Both gels could afterwards be re-crosslinked by red light exposure and displayed similar G’ values as in the initial hydrogel formation. Since 4-arm PEG-PhyB provided a superior dynamic range, with a more than 40-fold change of recorded stiffness, we chose this variant for further experiments.

**Figure 3:**
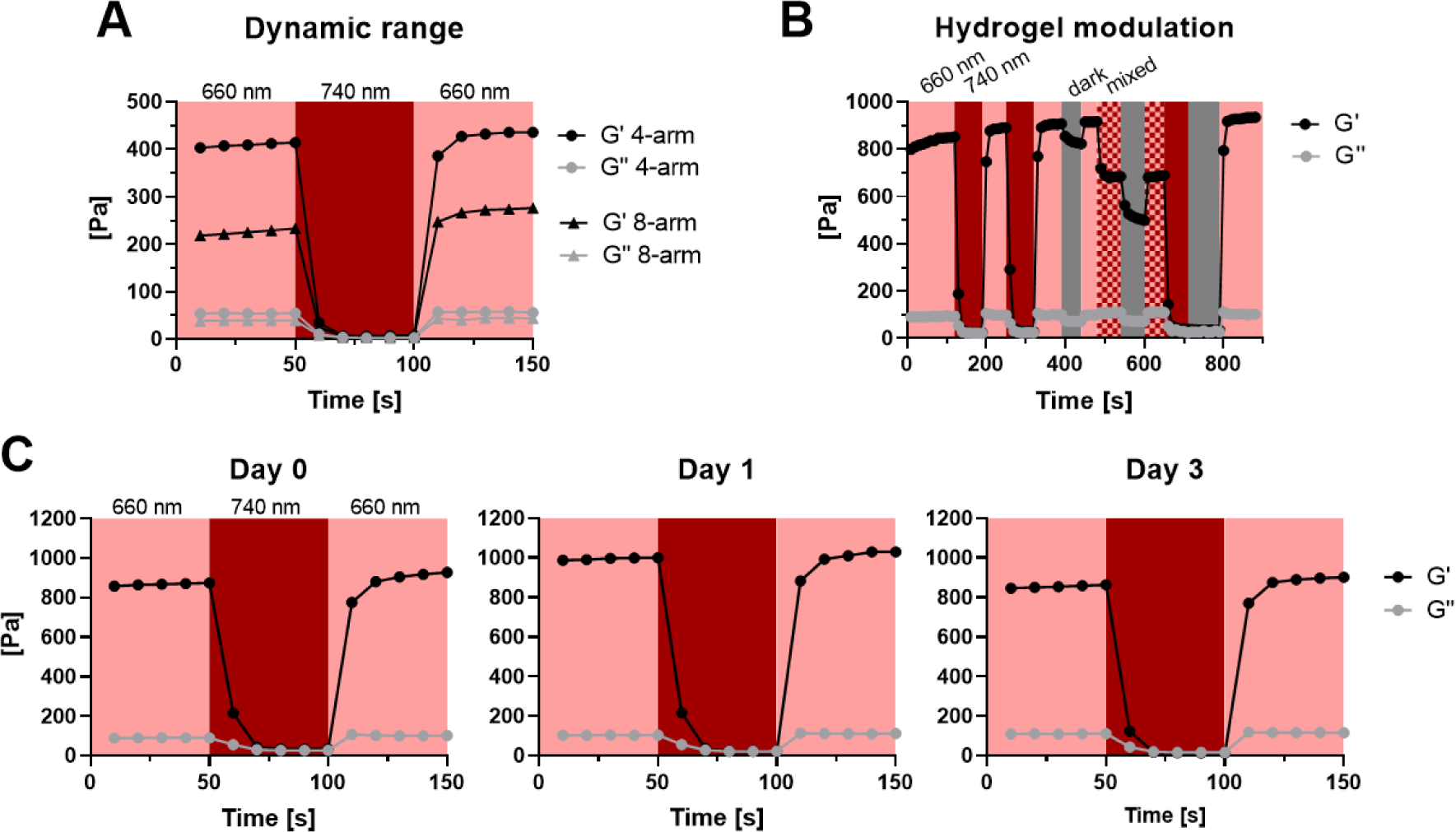
Characterization of light-dependent hydrogel mechanics. PEG-PhyB and PEG-PIF6 were mixed under 740 nm illumination and storage (G’) and loss (G’’) modulus of PEG-PhyB- and PEG-PIF6-based materials were recorded by small amplitude oscillatory shear rheology over time in response to different illumination conditions: 660 nm: light red; 740 nm: dark red; 660 and 740 nm light: checked; no light: grey. (A) 4-arm or 8-arm PEG-PhyB was mixed with 8-arm PEG-PIF6 (final coupled PhyB concentration: 50 mg/ml) under 740 nm illumination, illuminated with 660 nm for gelation and the light-dependent mechanical properties were recorded. (B) A hydrogel consisting of 4-arm PEG-PhyB and 8-arm PEG-PIF6 (final coupled PhyB concentration: 85 mg/ml) was prepared and analyzed as described in (A). (C) Dynamic range and functionality of the hydrogel variant from (B) was tested at 0, 24, and 72 h after initial hydrogel preparation.

Next, we further modified our formulation and tested whether an increase in protein concentration would correspond with a higher crosslinking density and as a result lead to a measurable increase in stiffness. In addition, we investigated hydrogel functionality after repeated illumination cycles including mixed light illumination (660 + 740 nm) and dark steps. Indeed, increasing the PEG-PhyB concentration to 85 mg/ml also raised the storage modulus to 832 ± 18 Pa (Figure 3B), demonstrating the tunability of gelation in response to the protein concentration. The light-adjustable stiffness of this hydrogel is in the range of soft biological tissues such as the brain and static materials with comparable stiffness have revealed important insight into cell-matrix interactions and mechanobiology.^[1,50–53]^ The light-dependent changes in stiffness were further shown to be completely reversible. Importantly, we were also able to achieve an intermediate stiffness of 683 ± 3 Pa, roughly 80% of the maximum, when both light conditions were applied simultaneously, demonstrating that the stiffness can be adjusted by tuning the colors and relative intensities of the applied light. Furthermore, the hydrogel demonstrated high functional stability over time by displaying an unaltered dynamic range.

We further demonstrated the functionality of the hydrogel for three days, a typical duration of cell culture experiments. To prevent microbial contamination, all reagents were sterile-filtered prior to use. In this context, we also confirmed compatibility of the hydrogel with chemicals typically added for cell culture experiments such as antibiotics (penicillin, streptomycin, and gentamycin) or protease inhibitors (optionally added to maintain hydrogel integrity).^[54,55]^ We investigated hydrogel stability and light responsiveness under these conditions by rheological measurements on day 0, 1, and 3 after initial hydrogel preparation (Figure 3C). At all time-points the hydrogels displayed similar G’ and G’’ values and demonstrated fully reversible gel-sol transitions.

Finally, we investigated the swelling ratio and stability of the crosslinked hydrogels in solution over time. For this purpose, we placed hydrogels in phosphate buffered saline (PBS) at room temperature (RT), illuminated them for 6 h with pulsed 660 nm light (5 min ON, 25 min OFF), and acquired at the beginning and after the end of each dark cycle an image (**Figure S5A**). We observed that the hydrogels remained intact for the entire experiment period and that the swelling reached a plateau after 6 h with a swelling ratio of 180 ± 17 % (Figure S5B). Importantly, this experiment also demonstrated that pulsed red light illumination is sufficient to retain structural integrity of the hydrogel, which is in agreement with the reported dark reversion half-life time of the used PhyB variant of t_1/2_ = 5.7 h.^[56]^

### 2.4 Hydrogel-controlled deposition of mammalian cells inside a microfluidic chip

Microfluidics refers to a range of technologies that allow working with small sample volumes inside a microstructure with sizes ranging from one to hundreds of microns and automating operations, that would normally require an entire laboratory.^[22]^ Their combination with *in-vitro* cell culture enables unprecedented user-defined control in research ranging from cell migration and cell-cell interaction studies to single cell genomics.^[57–62]^ By integrating our newly developed hydrogel into a microfluidic chip, we aimed to optically control the formation of geometrically confined hydrogels to encapsulate and subsequently deposit mammalian cells with high spatiotemporal precision inside the chip.

We first investigated the influence of the hydrogel on cell viability after encapsulation. To this aim, we synthesized hydrogels containing Chinese hamster ovary (CHO) cells at a concentration of 1 x 10^6^ cells/ml and incubated them under continuous 660 nm illumination for 1 h. As controls, we incubated adherent CHO cells for 1 h in cell culture medium or PBS (solvent of the hydrogel components) under 660 nm light or in the dark. After incubation, the cells were released from the hydrogels by illumination with 740 nm light, stained with a LIVE/DEAD marker (Zombie Violet Dye), and cell viability was assessed by flow cytometry (**Figure 4A**). We determined the cell viability after hydrogel encapsulation as 82 ± 3 % which was slightly lower compared to the viability of adherent CHO cells in PBS under red light illumination (95 ± 1 %, Figure 4B). In summary, these results indicate that after light-controlled cell encapsulation and subsequent release the vast majority of the cells is viable.

**Figure 4:**
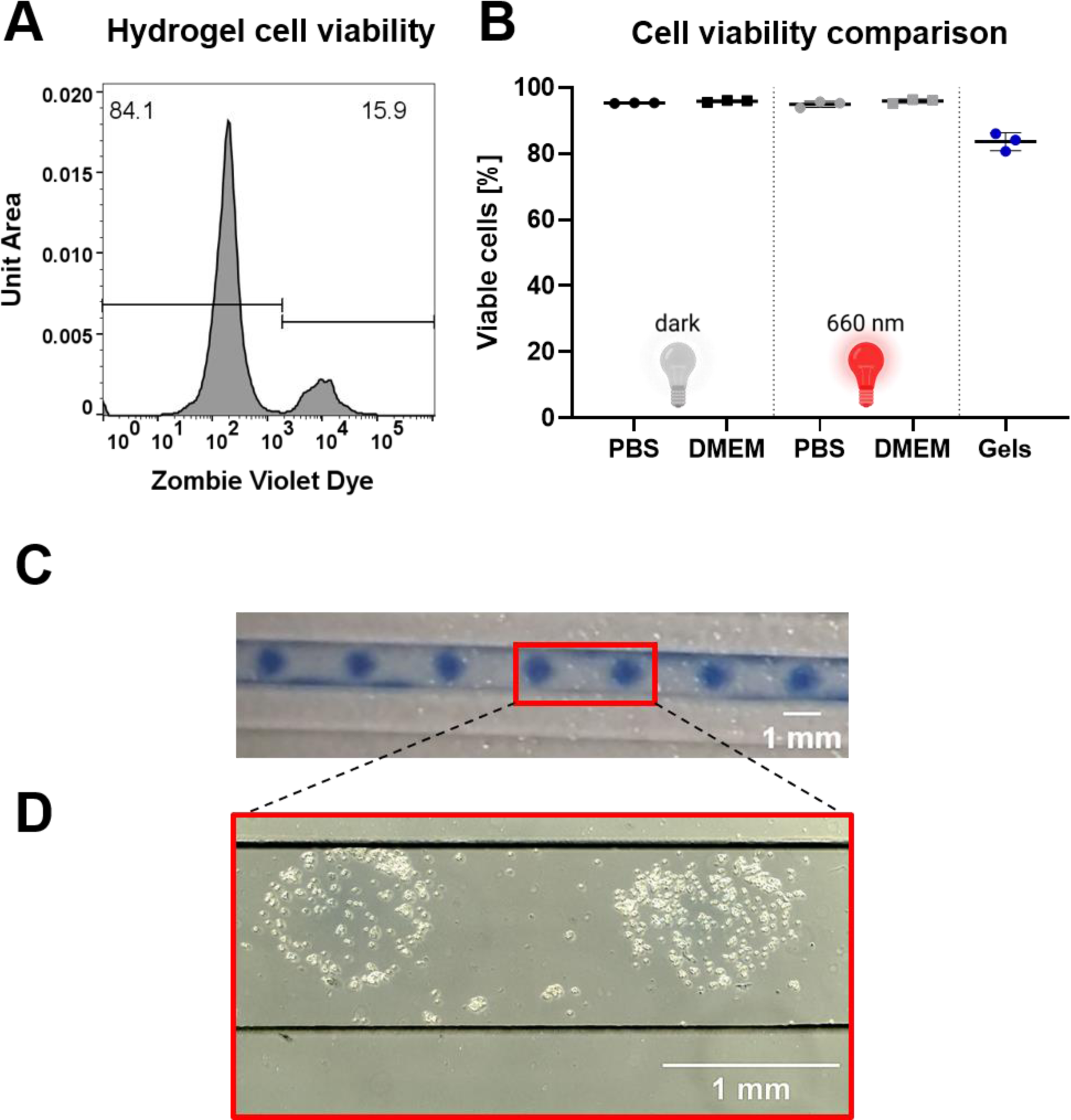
Light controlled spatially resolved deposition of cells within a microfluidic chip. (A) Viability of cells after encapsulation in hydrogel and subsequent release. 4-arm PEG-PhyB was mixed with 8-arm PEG-PIF6 (final coupled PhyB concentration: 50 mg/ml) and Chinese hamster ovary (CHO) cells (final concentration 1×10^6^ cells/ml) under 740 nm light. Subsequently, hydrogel formation was facilitated by 660 nm illumination for 1 h and cells were released afterwards by far-red illumination for 1 min. Cells were stained with a LIVE/DEAD marker and overall viability was assessed via flow cytometry. All cells were gated and LIVE/DEAD differentiation was performed as indicated in the graph. Area under curve was normalized (total area = 1) and histogram shown is representative of 3 samples. (B) Bar graph comparing cell viability after hydrogel encapsulation with different controls. For controls, CHO cells were incubated for 1 h in the dark or under 660 nm illumination overlaid with phosphate buffered saline (PBS) or cell culture medium. Afterwards, cells were stained and analyzed as described in (A). Individual values are shown, as well as mean ± SD (n = 3). (C) Spatially resolved hydrogel gelation in a microfluidic chip. Hydrogels as described in (A) were synthesized and CHO cells were added (final concentration: 150,000 cells/ml) under 740 nm light. Cell-polymer mixture was injected into a channel of a microfluidic chip and hydrogel formation was locally confined by patterned illumination with 660 nm light through a photomask. Finally, soluble components were removed by flushing the channel with PBS. (D) Deposited CHO cells within a microfluidic chip channel. Previously polymerized hydrogel from (C) was dissolved by illumination with far-red light (740 nm) to locally release the embedded mammalian cells.

For microfluidic cell deposition, CHO cells were mixed under far-red light illumination with 8-arm PEG-PIF6 and 4-arm PEG-PhyB (final coupled PhyB concentration: 50 mg/ml) to achieve a final concentration of 150,000 cells/ml. The cell-polymer mixture was then injected into a microfluidic channel of the chip and polymerization was initiated by local illumination with 660 nm light through a photomask to create localized hydrogels encapsulating the cells. Afterwards, the soluble material components were removed by flushing the channel with PBS and the hydrogel patterns produced by the photomask became clearly visible (Figure 4C). Finally, the embedded CHO cells were released by dissolving the gels via illumination with 740 nm light and microscopic images were taken (Figure 4D). These images revealed the successful confinement of the cells to the hydrogel areas and therefore demonstrated their light-guided positioning within a sub-millimeter geometry.

## 3. Conclusion

While stimuli-responsive hydrogels are a powerful technology to generate 3D matrices that resemble the dynamic, extracellular *in vivo* environment, they also present a beneficial combination with microfluidic technologies for various applications such as drug testing or organ-on-a-chip experiments.

In this work, we present an innovative optogenetic hydrogel whose mechanical properties can be reversibly controlled by illumination with cell-compatible, low intensity red/far-red light to adjust hydrogel stiffness from the sol state to ∼800 Pa, including intermediate states. Further, we harnessed the reversible sol-gel transition to place CHO cells with high spatiotemporal precision inside a microfluidic chip, opening the opportunity to optically program the architecture in situ within microfluidic devices. However, the high PhyB protein quantity required for hydrogel synthesis presents a limiting factor of this system, since considerable amounts are lost during the coupling and purification process.

The complete and reversible red light-dependent gel-to-sol transition enables unprecedented prospects for its application and displays a significant dynamic range with excellent functional stability. We expect the light-responsive hydrogel presented here to foster advances in the field of 3D cell culture, user-defined cell deposition, as well as microfluidics in biological applications.

## 4. Experimental Section

### 4.1 Cloning of expression plasmids

All expression plasmids (Figure S1) were assembled by Gibson cloning and the coding sequences of the different proteins are depicted in Table S1. For the PhyB expression plasmids pMH620 and pMH1414, the coding sequence of PhyB in plasmid pMH610^[63]^ was replaced by PhyB-Spy and PhyB-SpyΔ, respectively. The PhyB expression plasmids further encoded for the phycocyanobilin (PCB) biosynthesis enzymes heme oxygenase 1 (HO1) and PCB:ferredoxin oxidoreductase (PcyA) from *Synechocystis* PCC 6803. For the PIF6 expression plasmids pMH1411 and pMH613, the respective coding sequences were cloned into the vector pRSET (Invitrogen). The sequences of the open reading frames were verified by Sanger sequencing.

### 4.2 Protein production

For PhyB-Spy protein production the plasmid (pMH620) was transformed into *E. coli* strain BL21 Star (DE3) (Thermo Fisher Scientific) and grown overnight at 37°C in the presence of streptomycin (100 µg/ml). On the next day, expression culture was inoculated with this pre-culture and grown at 30°C until an OD_600_ of 0.6 – 0.8 was reached. At this point, isopropyl-β-D-thiogalactopyranoside (IPTG, 1 mM) was added to induce protein production and bacteria were cultured at 18°C for 20-24 h in the dark. Cells were harvested by centrifugation (8 min, 6500 g) and cell pellets were resuspended in Ni-Lysis buffer (50 mM NaH_2_PO_4_, 300 mM NaCl, 10 mM imidazole, pH 8.0) before being shock frozen in liquid nitrogen and stored at −80°C. PhyB-SpyΔ (pMH1414) protein was produced by high-cell-density *E. coli* fermentation as described previously.^[63]^

For PIF6 and moxGFP-PIF6 production, BL21(DE3) pLysS *E. coli* cells (Thermo Fisher Scientific) were transformed with plasmid pMH613 or pMH1411, respectively, and grown overnight at 37°C in the presence of ampicillin (100 µg/ml) and chloramphenicol (34 µg/ml). Afterwards, proteins were produced at 25 °C overnight as described for PhyB-Spy.

### 4.3 Protein purification

Frozen cell pellets were thawed at 37 °C in a water bath, resuspended in Ni-Lysis buffer followed by cell lysis using a high-pressure homogenizer (1200 bar, model: APV-2000, SPX Flow Technology). The lysate was clarified by centrifugation (1 h, 30,000 g) and the clarified lysate subjected to immobilized metal affinity chromatography (IMAC). To this end, the clarified lysate was loaded on a Ni-nitrilotriacetic acid (NTA) agarose column (Qiagen) using the ÄKTA Explorer fast protein liquid chromatography system (FPLC, GE Healthcare). Afterwards, the column was washed with 12 column volumes (CV) of Ni-wash buffer (50 mM NaH_2_PO_4_, 300 mM NaCl, 20 mM imidazole, pH 8.0) followed by protein elution with 6 CV of Ni-elution buffer (50 mM NaH_2_PO_4_, 300 mM NaCl, 250 mM imidazole, pH 8.0).

### 4.4 Protein characterization

For SDS-PAGE analysis, proteins were incubated with 1x SDS loading buffer (from 5x stock solution with 50% (v/v) Glycerol, 312.5 mM Tris, 12.5% (v/v) 2-mercaptoethanol (2-ME), 10% (w/v) SDS, 0.05% Bromophenol blue) at 95°C for 5 min. After gel electrophoresis, the gel was incubated in a 1 mM zinc acetate solution for 15 min before visualization of zinc-induced chromophore fluorescence using UV light (312 nm). Afterwards, proteins on the gels were stained with Coomassie solution. The concentrations of the proteins were determined photometrically by measuring the absorbance at 280 nm or by Bradford assay using bovine serum albumin (BSA) as standard.

### 4.5 Coupling of PhyB to PEG by Spy Chemistry

First, SpyTag peptide (N-term-AHIVMVDAYKPTKGSGDRCG-C-term, N-term: Acetylation, C-term: Amidation, purity: 94%, GenScript Biotech) was dissolved in triethanolamine (TEA, 0.3 M), mixed with 8-arm PEG-VS (40 kDa, NOF Europe, cat. no. Sunbright HGEO-400VS) or 4-arm PEG-VS (20 kDa, JenKem Technology USA, cat. no. A7025-1) in a peptide:VS molar ratio of 1.5:1, and incubated overnight at room temperature (RT). Afterwards, uncoupled VS-groups were blocked by incubation with 2-ME (10 mM) for 30 min at RT and uncoupled peptide was removed by dialysis (10 kDa MWCO) against water for 2 days. Next, the conjugated PEG was lyophilized and dissolved again in phosphate buffered saline (PBS, (2.7 mM KCl, 1.5 mM KH_2_PO_4_, 8.1 mM Na_2_HPO_4_, 137 mM NaCl) at a concentration of 5% (w/v). Finally, PEG-SpyTag was incubated with purified PhyB (∼ 30 mg/ml) for 2 h at RT.

### 4.6 Coupling of PIF6 to PEG by Michael-type addition

First, the elution buffer of purified PIF6 was changed to reaction buffer (1x PBS, 2 mM EDTA, pH 8.0) using a desalting column (5 kDa MWCO) and the protein was concentrated by ultrafiltration (10 kDa MWCO PES) to a concentration of ∼ 45 mg/ml. Next, the protein was reduced for 1 h at room temperature with TCEP (from 100 mM stock in 0.5 M NaHCO_3_, pH 8.0) in a molar TCEP:protein ratio of 0.7:1 and mixed at a concentration of ∼ 30 mg/ml with 8-arm PEG-VS (molar VS:protein ratio of 1:3) in reaction buffer with TEA (final concentration of 100 mM from 1 M stock in reaction buffer, pH 8.0). After incubation overnight at RT, 2-ME (10 mM) was added and incubated for 30 min to block uncoupled VS-groups.

### 4.7 Size-exclusion chromatography (SEC)

Coupled PEG-PhyB and PEG-PIF6 were purified by size exclusion chromatography using a HiLoad Superdex 200 pg (16/600 or 25/600, GE Healthcare) column connected to an ÄKTA Explorer system with PBS as a running buffer. Analytical SEC was performed with a Superdex 200 Increase 10/300 GL (GE Healthcare) column using PBS as a running buffer. For the analysis of the light-dependent protein interaction between PhyB and PIF6, samples were illuminated for 2 min with 660 or 740 nm light (100 μmol/m^2^/s) and the subsequent SEC was performed in the dark.

### 4.8 Hydrogel synthesis, rheological measurements and swelling analysis

Purified PEG-PhyB and PEG-PIF6 were concentrated by ultrafiltration and mixed at the indicated final coupled PhyB concentration (molar PhyB:PIF6 ratio of 1:2) under 740 nm light (100 μmol/m^2^/s). Hydrogels were characterized by small amplitude oscillatory shear experiments using an MCR301 rheometer (Anton Paar) with parallel plate configuration. To this aim, 25 µl of the hydrogel solution was pipetted under 740 nm light on the lower glass plate (P-PTD 120, Anton Paar) and after illumination with 660 nm (270 μmol/m^2^/s) for 2 min, the hydrogel was compressed with the upper measuring plate (8 mm diameter, PP08, Anton Paar) until a gap size of 250 µm was reached. Measurements were performed with 1 Hz frequency and 2% strain. To prevent dehydration of the hydrogels during measurements, silicone oil was added around the gel. Illumination was performed through the lower glass plate using 660 nm (270 μmol/m^2^/s) and 740 nm (180 μmol/m^2^/s) LEDs. For swelling analysis 15 µl of the hydrogel mix were added on siliconized (Sigmacote, Sigma, cat. no. SL-2) glass slides. Hydrogels were then crosslinked for 1 min by illumination at 660 nm (40 μmol/m^2^/s) and transferred into 500 µl PBS inside a 48-well plate. A reference image was retrieved immediately after, while placing the plate above grid paper as scale bar. Gels were then subjected to illumination cycles consisting of 5 min red light (40 μmol/m^2^/s) illumination, followed by a 25 min dark period. Gel size was analyzed by referencing a 5 mm scale-bar to measure the horizontal diameter of the hydrogels in triplicates. Volumetric swelling was calculated assuming isotropic swelling from the reference image at timepoint zero.

### 4.9 Cell viability

CHO cells were cultured in cell culture medium (DMEM supplemented with 10% FCS, 100 U/ml penicillin and 100 µg/ml streptomycin) at 37°C in a humidified atmosphere with 5% CO_2_. For analysis of cell viability, cells were detached using trypsin, centrifuged (5 min, 300 g) and resuspended in PBS. Subsequently, cells were added to 30 µl hydrogel (final coupled PhyB concentration of 50 mg/ml) under 740 nm far-red light illumination to a final concentration of 1 x 10^6^ cells/ml. Samples were subsequently illuminated with 660 nm red light (40 μmol/m^2^/s) at RT for 1 hour to facilitate hydrogel formation. Afterwards, hydrogels were dissolved by far-red light exposure (100 μmol/m^2^/s) and cells were subjected to viability staining. To this aim, the dissolved hydrogel was transferred into a V-bottom 96-well plate and subsequently centrifuged (4 min, 500 g). Cells were then washed by resuspending in 200 µl PBS, centrifuged (4 min, 500 g), resuspended in 50 µl LIVE/DEAD staining solution (PBS with 1:500 Zombie Violet Dye, Biolegend cat. no. 423113) and incubated for 20 min at 4 °C in the dark. After incubation, cells were washed with PBS and resuspended in 200 µl buffer (PBS supplemented with 2% FCS) and analyzed via Attune NxT Flow Cytometer (ThermoFisher). For LIVE/DEAD analysis all cells were gated and viable cells were quantified as indicated in Figure 4 via FlowJo (Version 10.9.0) software. For controls, medium was removed from CHO cells cultured in a 6-well plate and 2 ml PBS or cell culture medium were added before incubation for 1 h at RT with 660 nm illumination (40 μmol/m^2^/s) or in the dark. Afterwards, supernatant medium was collected in 5 ml DMEM and cells were detached using trypsin and pooled with the collected supernatant. Cells were centrifuged (4 min, 500 g) and resuspended in 1 ml PBS. Cell number was determined and 30,000 cells were transferred into a 96-well V-bottom plate for subsequent staining and analysis as described above.

### 4.10 Deposition of mammalian cells in microfluidic chip

CHO cells were cultured and detached as described above. For cell deposition experiments, cells were resuspended in cell culture medium. Cells were then mixed with hydrogel solution (final coupled PhyB concentration of 50 mg/ml, molar PhyB:PIF6 ratio of 1:1.5) under 740 nm light (100 μmol/m^2^/s) at a final concentration of 150,000 cells/ml and 20 µl were injected into a channel of a microfluidic chip. Afterwards, the channel was illuminated for 3 min in a spatially restricted manner with 660 nm light (20 μmol/m^2^/s) through a photomask and the non-gelated material was removed by flushing the channel with 200 µl PBS. Afterwards, the cell embedding hydrogel was dissolved by illumination with 740 nm light (100 μmol/m^2^/s) for 2 min and the CHO cells were visualized by microscopy.

### Statistics

If not stated otherwise, the values indicated in the main text represent mean ± SD. For rheological characterization, only the plateau phase of the first illumination cycle of the corresponding condition was analyzed.

## Figures

Figures 1, 2 and S1 were created with BioRender.com.

## Supporting Information

Supporting Information is available from the Wiley Online Library or from the author.

## Data availability statement

The data that support the findings of this study are available from the corresponding author upon reasonable request.

## Acknowledgements

We thank the technical workshop of the Faculty of Biology for the design and construction of illumination devices and acknowledge the scientific assistance of the Core Facility Signalling Factory for their assistance with size exclusion experiments. This work was supported by the European Union (ERC, STEADY, 101053857), the German Research Foundation (Deutsche Forschungsgemeinschaft, DFG) under Germany’s Excellence Strategy – CIBSS, EXC-2189, Project ID: 390939984 and under the Excellence Initiative of the German Federal and State Governments – BIOSS, EXC-294, and in part by the Ministry for Science, Research and Arts of the State of Baden-Württemberg.

