## Supporting information for "A photoreceptor-based hydrogel with red light-responsive reversible sol-gel transition as transient cellular matrix"

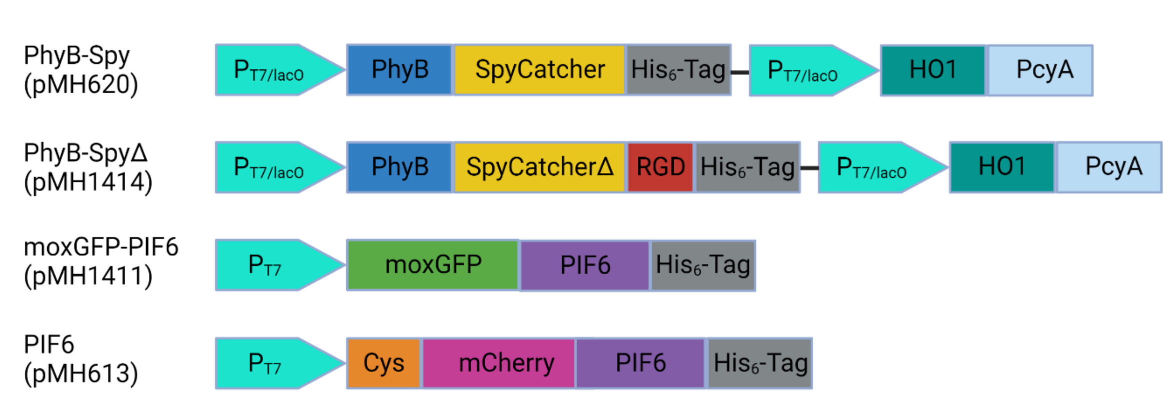


**Figure S 1: Overview of plasmids**

Schematic presentation of the promotor and open reading frames including $His_6$-Tag for purification, Cys motif (GCRD) for Michael-type addition and RGD sequence for cell adhesion.

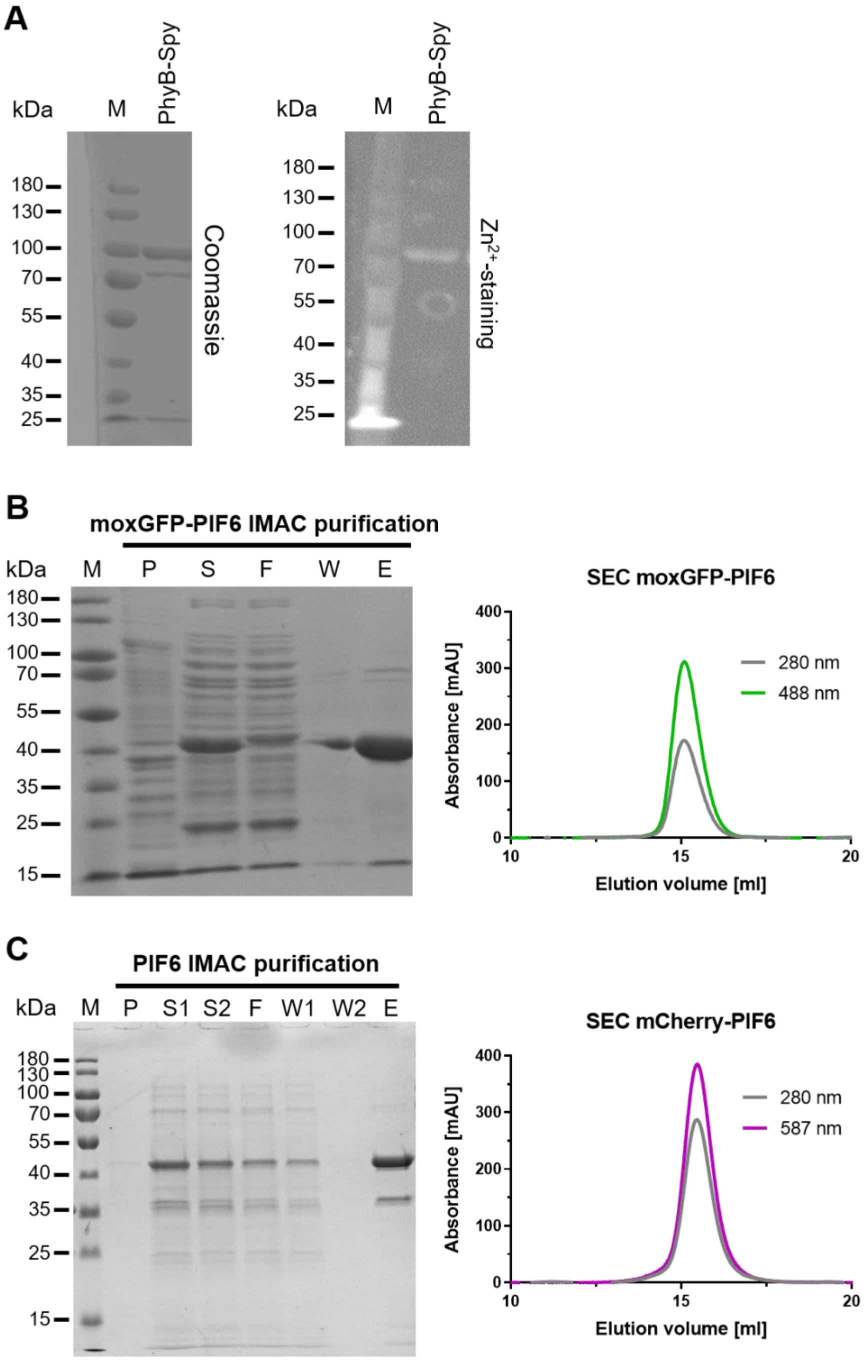


**Figure S 2: Characterization of purified PhyB-Spy, moxGFP-PIF6 and PIF6.**

(A) SDS PAGE followed by Coomassie and $Zn^{2+}$ staining of IMAC purified PhyB-Spy (pMH620). Purified protein displayed the expected size of 85 kDa.

(B + C) SDS-PAGE (left) and SEC (right) analysis of IMAC purified moxGFP-PIF6 (B, pMH1411) and mCherry-PIF6 (C, pMH613, referred to as PIF6) after recombinant protein expression in *E. coli*. Fractions of the purification process were collected and visualized by SDS-PAGE and Coomassie staining (M: molecular ladder, P: insoluble fraction of *E. coli* lysate; S: soluble fraction of lysate; F: flow-through of IMAC column; W: wash of column; E: eluate of column). Eluted proteins correlated with expected size of 39 kDa (B) or 40 kDa (C). Purity was further assessed by SEC and monitoring moxGFP-PIF6 (488 nm) or PIF6 (587 nm) specific absorbance as well as protein absorbance (280 nm).

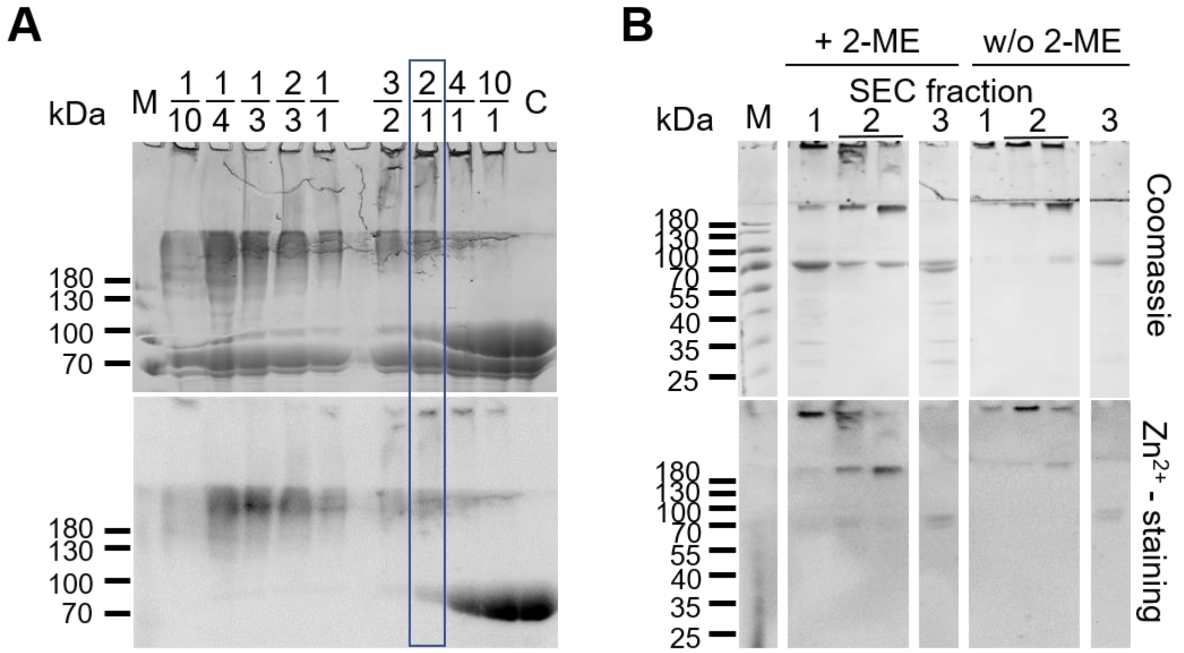


**Figure S 3: Determination of the optimal ratio of PhyB-SpyCatcher coupling to PEG-SpyTag and analysis of the fractions of the subsequent SEC for purification.**

(A) PhyB-Spy and SpyTag functionalized PEG were incubated in protein:polymer ratios ranging from 0.1 to 10 as indicated in the picture. Afterwards, coupling efficiency was determined by SDS-PAGE, followed by Coomassie and $Zn^{2+}$ staining (M: molecular ladder; C: uncoupled PhyB-Spy protein). The optimal ratio with minimal remaining uncoupled protein was selected as 2:1 (highlighted with blue frame). (B) Pooled fractions of the indicated areas in Figure 2C (1, 2, and 3) were analyzed by SDS-PAGE and visualized by Coomassie and $Zn^{2+}$ staining (expected MW of uncoupled PhyB-Spy: 85 kDa), either with (+2-ME) or without (w/o 2-ME) 2-ME (12.5%) in the loading buffer (M: molecular ladder).

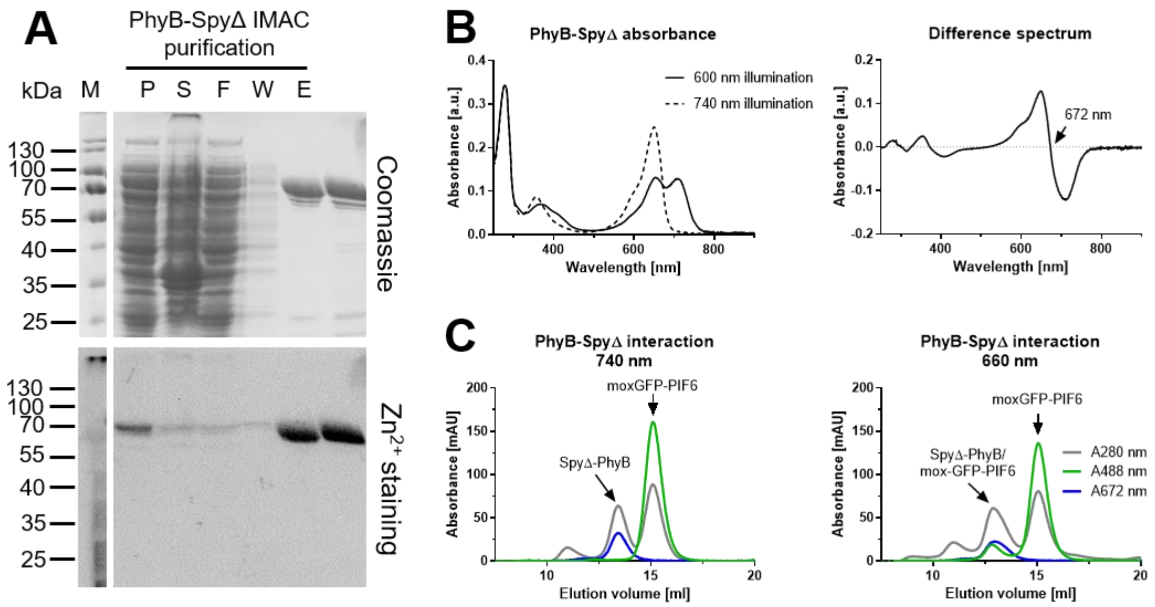


**Figure S 4: Purification and characterization PhyB-SpyΔ.**

(A) PhyB-SpyΔ was purified by IMAC, characterized by SDS PAGE and protein bands visualized by Coomassie and $Zn^{2+}$ staining (M: molecular ladder, P: insoluble fraction of *E. coli* lysate; S: soluble fraction of lysate; F: flow-through of IMAC column; W: wash of column; E: eluate of column). Eluted protein had an expected size of 83 kDa and was compared to the original PhyB-Spy purified protein. (B) Absorbance spectrum of PhyB-SpyΔ after illumination with red (660 nm) or far-red (740 nm) light. Difference spectrum of both conditions was calculated and the isosbestic point determined at 672 nm. (C) PhyB-SpyΔ was mixed with moxGFP-PIF6 after illumination with red or far-red light. Afterwards, protein interaction was analyzed by SEC and monitoring protein absorbance at 280 nm as well as PhyB (672 nm) and moxGFP-PIF6 (488 nm) specific absorbance. a.u. = arbitrary unit

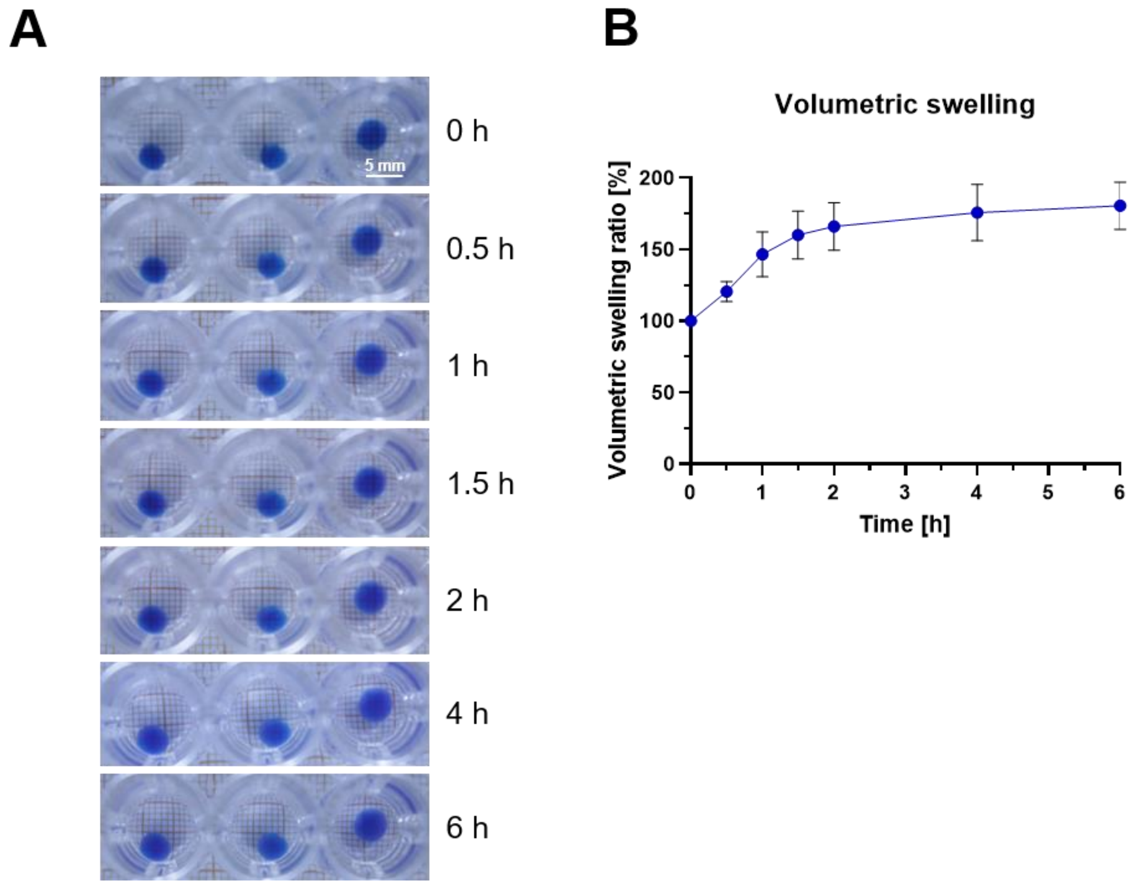


**Figure S 5: Stability and swelling of hydrogels in solution over time.**
(A) Hydrogels were synthesized by mixing 4-arm PEG-PhyB (final coupled PhyB concentration: 50 mg/ml) with 8-arm PEG-PIF6 under far-red (740 nm) light. Pipetting 15 µl of the mixture onto a siliconized glass slide was followed by crosslinking with red (660 nm) light for 1 min. Afterwards, the hydrogels were transferred into wells of a 48-well plate containing 500 µl PBS and illuminated with pulsed 660 nm light (5 min ON, 25 min OFF). After the end of each dark cycle, an image was acquired. (B) Horizontal hydrogel diameter was measured in triplicates from images retrieved in (A) using a grid paper and the volumetric swelling was calculated assuming isotropic swelling. Values are shown as mean ± SD (n = 3).

**Table S1: Plasmid sequences.**
Left column shows encoded protein, internal designated plasmid number (pMH) and a color code for the functional areas of the plasmid. Right side displays open reading frames of plasmids as well as promotor region. Uncolored sequences are promotor regions or linker sequences.

| Construct | Sequence |
|---|---|
| **PhyB-SpyCatcher (pMH620)** | ...ATTAGGAAATTAATACGACTCACTATAGGGGAATTGTGAGCGGATAACAATTCCCCTGTAGAAA<br>TAATTTTGTTTAACTTTAATAAGGAGATATACCATGGTTAGCGGTGTTGGTGGTAGCGGTGGTGGT<br>CGTGGTGGCGGTCGCGGAGGTGAAGAAGAACCGAGCAGCAGCCATACCCCGAATAATCGCCGT<br>GGTGGTGAACAGGCACAGAGCAGCGGCACCAAAAGCCTGCGTCCGCGTAGCAATACCGAAAGC |

T7 promotor
PhyB (1-651)
SpyCatcher
$His_6$-Tag
HO1
PcyA
Backbone: pCDFDuet-1 (Novagen)

ATGAGCAAAGCAATTCAGCAGTATACCGTTGATGCACGTCTGCATGCCGTTTTCGAACAGAGCGG
TGAAAGCGGTAAAAGCTTTGATTATAGCCAGAGCCTGAAAACCACCACCTATGGTAGCAGCGTGC
CGGAACAGCAGATTACCGCATATCTGAGCCGTATTCAGCGTGGTGGCTACATTCAGCCGTTTGG
CTGCATGATCGCAGTTGATGAAAGCAGCTTTCGCATTATTGGCTACAGCGAAAATGCACGTGAAA
TGCTGGGCATTATGCCGCAGAGCGTTCCGACCCTGGAAAAACCGGAAATTCTGGCAATGGGCAC
CGATGTTCGTAGCCTGTTTACCAGCAGCAGCTCCATTCTGCTGGAACGTGCCTTTGTTGCCCGTG
AAATTACCCTGCTGAATCCGGTTTGGATTCATAGCAAAAACACCGGCAAACCGTTTTATGCAATTC
TGCATCGTATTGATGTTGGCGTGGTTATTGATCTGGAACCGGCACGTACCGAAGATCCGGCACT
GAGCATTGCCGGTGCAGTTCAGAGCCAGAAACTGGCAGTTCGTGCAATTAGCCAGCTGCAGGCA
CTGCCTGGTGGTGATATCAAACTGCTGTGTGATACCGTTGTTGAAAGCGTTCGTGATCTGACCGG
CTACGACCGTGTTATGGTGTATAAATTCCACGAAGATGAACATGGTGAAGTTGTTGCAGAAAGCA
AACGTGATGACCTGGAACCGTATATTGGTCTGCATTATCCAGCAACCGATATTCCGCAGGCAAGC
CGTTTCCTGTTCAAACAGAATCGTGTGCGCATGATTGTTGATTGTAATGCAACACCGGTTCTGGTT
GTTCAGGATGATCGTCTGACCCAGAGCATGTGTCTGGTTGGTAGCACCCTGCGTGCACCGCATG
GTTGTCATAGCCAGTATATGGCAAATATGGGTAGCATCGCAAGCCTGGCCATGGCGGTGATCAT
CAATGGTAATGAAGATGATGGTAGCAATGTTGCAAGCGGTCGTAGCAGCATGCGTCTGTGGGGT
CTGGTTGTGTGTCATCATACCAGCAGTCGCTGCATTCCGTTTCCGCTGCGTTATGCATGTGAATT
TCTGATGCAGGCATTTGGACTGCAGCTGAATATGGAACTGCAACTGGCACTGCAGATGAGCGAA
AAACGTGTTCTGCGTACCCAGACCCTGCTGTGCGATATGCTGCTGCGTGATAGTCCGGCAGGCA
TTGTTACCCAGAGCCCGAGCATTATGGATCTGGTGAAATGCGATGGTGCAGCCTTTCTGTATCAC
GGTAAATACTATCCGCTGGGTGTTGCACCGAGCGAAGTTCAGATTAAAGATGTTGTTGAGTGGCT
GCTGGCAAATCATGCAGATAGCACCGGTCTGAGCACCGATAGCCTGGGTGATGCAGGTTATCCG
GGTGCAGCAGCACTGGGAGATGCAGTTTGTGGTATGGCAGTTGCATACATTACCAAACGCGATT
TTCTGTTTTGGTTTCGTAGCCATACCGCCAAAGAAATCAAATGGGGTGGTGCAAAACATCACCCG
GAAGATAAAGATGACGGTCAGCGTATGCATCCGCGTAGTAGCTTTCAGGCATTTCTGGAAGTGGT
GAAAAGCCGTAGCCAGCCGTGGGAAACCGCAGAAATGGATGCAATTCATAGCCTGCAACTGATT
CTGCGCGATAGCTTCAAAGAAAGCGAAGCAGCAATGAATAGCAAAGTTGTTGATGGTGTTGTTCA
GCCGTGTCGTGATATGGCAGGCGAACAGGGTATTGATGAACTGGGTGCAGGTTCTGGTAGCGGT
GGCGCAATGGTTGATACCTTATCAGGTTTATCAAGTGAGCAAGGTCAGTCCGGTGATATGACAAT
TGAAGAAGATAGTGCTACCCATATTAAATTCTCAAAACGTGATGAGGACGGCAAAGAGTTAGCTG
GTGCAACTATGGAGTTGCGTGATTCATCTGGTAAAACTATTAGTACATGGATTTCAGATGGACAA
GTGAAAGATTTCTACCTGTATCCAGGAAAATATACATTTGTCGAAACCGCAGCACCAGACGGTTA
TGAGGTAGCAACTGCTATTACCTTTACAGTTAATGAGCAAGGTCAGGTTACTGTAAATGGCAAAG
CAACTAAAGGTGACGCTCATATTCATCATCACCATCACCATTAAGCGGCCGCATAATGCTTAAGT
CGAACAGAAAGTAATCGTATTGTACACGGCCGCATAATCGAAATTAATACGACTCACTATAGGGG
AATTGTGAGCGGATAACAATTCCCCATCTTAGTATATTAGTTAAGTATAAGAAGGAGATATACATA
TGAGTGTCAACTTAGCTTCCCAGTTGCGGGAAGGGACGAAAAAATCCCACTCCATGGCGGAGAA
CGTCGGCTTTGTCAAATGCTTCCTCAAGGGCGTTGTCGAGAAAAATTCCTACCGTAAGCTGGTTG
GCAATCTCTACTTTGTCTACAGTGCCATGGAAGAGGAAATGGCAAAATTTAAGGACCATCCCATC
CTCAGCCACATTTACTTCCCCGAACTCAACCGCAAACAAAGCCTAGAGCAAGACCTGCAATTCTA
TTACGGCTCCAACTGGCGGCAAGAAGTGAAAATTTCTGCCGCTGGCCAAGCCTATGTGGACCGA
GTCCGGCAAGTGGCCGCTACGGCCCCTGAATTGTTGGTGGCCCATTCCTACACCCGTTACCTGG
GGGATCTTTCCGGCGGTCAAATTCTCAAGAAAATTGCCCAAAATGCCATGAATCTCCACGATGGT
GGCACAGCTTTCTATGAATTTGCCGACATTGATGACGAAAAGGCTTTTAAAAATACCTACCGTCAA
GCTATGAATGATCTGCCCATTGACCAAGCCACCGCCGAACGGATTGTGGATGAAGCCAATGACG
CCTTTGCCATGAACATGAAAATGTTCAACGAACTTGAAGGCAACCTGATCAAGGCGATCGGCATT
ATGGTGTTCAACAGCCTCACCCGTCGCCGCAGTCAAGGCAGCACCGAAGTTGGCCTCGCCACCT
CCGAAGGCTAGAAGCTTGGATCCATTAAAGAGGAGAAACCCGGGCATGGCCGTCACTGATTTAA
GTTTGACCAATTCTTCCCTGATGCCTACGTTGAACCCGATGATTCAACAGTTGGCCCTGGCGATC
GCCGCTAGTTGGCAAAGTTTACCCCTCAAGCCCTATCAATTGCCGGAGGATTTGGGCTACGTAG
AAGGCCGCCTGGAAGGGGAAAAGTTAGTGATTGAAAATCGGTGCTACCAAACGCCCCAGTTTCG

<table>
<tr><td></td><td>CAAAATGCATTTGGAGTTGGCCAAGGTGGGCAAAGGGTTGGATATTCTCCACTGTGTAATGTTTC<br>CTGAGCCTTTATACGGTCTACCTTTGTTTGGCTGTGACATTGTGGCCGGCCCCGGTGGAGTAAGT<br>GCGGCTATTGCGGATCTATCCCCCACCCAAAGCGATCGCCAATTGCCCGCAGCGTACCAAAAAT<br>CATTGGCAGAGCTAGGCCAGCCAGAATTTGAGCAACAACGGGAATTGCCCCCCTGGGGAGAAAT<br>ATTTTCTGAATATTGTTTATTCATCCGTCCCAGCAATGTCACTGAAGAAGAAAGATTTGTACAAAG<br>GGTAGTGGACTTTTTGCAAATTCATTGTCACCAATCCATCGTTGCCGAACCCTTGTCTGAAGCTCA<br>AACTTTGGAGCACCGTCAGGGGCAAATTCATTACTGCCAACAACAACAGAAAAATGATAAAACCC<br>GTCGGGTACTGGAAAAAGCTTTTGGGGAAGCTTGGGCGGAACGGTATATGAGCCAAGTCTTATT<br>TGATGTTATCCAATAAGACGTCGGTACCCTCGAGTCTGGTAAAGAAACCGCTGCTGCGAAATTTG<br>AACGCCA…</td></tr>
<tr><td>PhyB-SpyΔ (pMH1414)<br><br>T7 promotor<br>PhyB (1-651)<br>SpyCatcher<br>RGD<br>His6-Tag<br>HO1<br>PcyA<br>Backbone: pCDFDuet-1 (Novagen)</td><td>…ATTAGGAAATTAATACGACTCACTATAGGGGAATTGTGAGCGGATAACAATTCCCCTGTAGAAA<br>TAATTTTGTTTAACTTTAATAAGGAGATATACCATGGTTAGCGGTGTTGGTGGTAGCGGTGGTGGT<br>CGTGGTGGCGGTCGCGGAGGTGAAGAAGAACCGAGCAGCAGCCATACCCCGAATAATCGCCGT<br>GGTGGTGAACAGGCACAGAGCAGCGGCACCAAAAGCCTGCGTCCGCGTAGCAATACCGAAAGC<br>ATGAGCAAAGCAATTCAGCAGTATACCGTTGATGCACGTCTGCATGCCGTTTTCGAACAGAGCGG<br>TGAAAGCGGTAAAAGCTTTGATTATAGCCAGAGCCTGAAAACCACCACCTATGGTAGCAGCGTGC<br>CGGAACAGCAGATTACCGCATATCTGAGCCGTATTCAGCGTGGTGGCTACATTCAGCCGTTTGG<br>CTGCATGATCGCAGTTGATGAAAGCAGCTTTCGCATTATTGGCTACAGCGAAAATGCACGTGAAA<br>TGCTGGGCATTATGCCGCAGAGCGTTCCGACCCTGGAAAAACCGGAAATTCTGGCAATGGGCAC<br>CGATGTTCGTAGCCTGTTTACCAGCAGCAGCTCCATTCTGCTGGAACGTGCCTTTGTTGCCCGTG<br>AAATTACCCTGCTGAATCCGGTTTGGATTCATAGCAAAAACACCGGCAAACCGTTTTATGCAATTC<br>TGCATCGTATTGATGTTGGCGTGGTTATTGATCTGGAACCGGCACGTACCGAAGATCCGGCACT<br>GAGCATTGCCGGTGCAGTTCAGAGCCAGAAACTGGCAGTTCGTGCAATTAGCCAGCTGCAGGCA<br>CTGCCTGGTGGTGATATCAAACTGCTGTGTGATACCGTTGTTGAAAGCGTTCGTGATCTGACCGG<br>CTACGACCGTGTTATGGTGTATAAATTCCACGAAGATGAACATGGTGAAGTTGTTGCAGAAAGCA<br>AACGTGATGACCTGGAACCGTATATTGGTCTGCATTATCCAGCAACCGATATTCCGCAGGCAAGC<br>CGTTTCCTGTTCAAACAGAATCGTGTGCGCATGATTGTTGATTGTAATGCAACACCGGTTCTGGTT<br>GTTCAGGATGATCGTCTGACCCAGAGCATGTGTCTGGTTGGTAGCACCCTGCGTGCACCGCATG<br>GTTGTCATAGCCAGTATATGGCAAATATGGGTAGCATCGCAAGCCTGGCCATGGCGGTGATCAT<br>CAATGGTAATGAAGATGATGGTAGCAATGTTGCAAGCGGTCGTAGCAGCATGCGTCTGTGGGGT<br>CTGGTTGTGTGTCATCATACCAGCAGTCGCTGCATTCCGTTTCCGCTGCGTTATGCATGTGAATT<br>TCTGATGCAGGCATTTGGACTGCAGCTGAATATGGAACTGCAACTGGCACTGCAGATGAGCGAA<br>AAACGTGTTCTGCGTACCCAGACCCTGCTGTGCGATATGCTGCTGCGTGATAGTCCGGCAGGCA<br>TTGTTACCCAGAGCCCGAGCATTATGGATCTGGTGAAATGCGATGGTGCAGCCTTTCTGTATCAC<br>GGTAAATACTATCCGCTGGGTGTTGCACCGAGCGAAGTTCAGATTAAAGATGTTGTTGAGTGGCT<br>GCTGGCAAATCATGCAGATAGCACCGGTCTGAGCACCGATAGCCTGGGTGATGCAGGTTATCCG<br>GGTGCAGCAGCACTGGGAGATGCAGTTTGTGGTATGGCAGTTGCATACATTACCAAACGCGATT<br>TTCTGTTTTGGTTTCGTAGCCATACCGCCAAAGAAATCAAATGGGGTGGTGCAAAACATCACCCG<br>GAAGATAAAGATGACGGTCAGCGTATGCATCCGCGTAGTAGCTTTCAGGCATTTCTGGAAGTGGT<br>GAAAAGCCGTAGCCAGCCGTGGGAAACCGCAGAAATGGATGCAATTCATAGCCTGCAACTGATT<br>CTGCGCGATAGCTTCAAAGAAAGCGAAGCAGCAATGAATAGCAAAGTTGTTGATGGTGTTGTTCA<br>GCCGTGTCGTGATATGGCAGGCGAACAGGGTATTGATGAACTGGGTGCAGGTTCTGGTAGCGGT<br>GGCGCAAGCGGTGATAGTGCTACCCATATTAAATTCTCAAAACGTGATGAGGACGGCAAAGAGTT<br>AGCTGGTGCAACTATGGAGTTGCGTGATTCATCTGGTAAAACTATTAGTACATGGATTTCAGATG<br>GACAAGTGAAAGATTTCTACCTGTATCCAGGAAAATATACATTTGTCGAAACCGCAGCACCAGAC<br>GGTTATGAGGTAGCAACTGCTATTACCTTTACAGTTAATGAGCAAGGTCAGGTTACTGTAAATGG<br>CAGCGGTGCTGGCTCCGCAGGTTCTGGCCGTGGCGATAGCCCAGGTCGTGGTGACTCTCCGCA<br>TCATCACCATCACCATTGAGCGGCCGCATAATGCTTAAGTCGAACAGAAAGTAATCGTATTGTAC<br>ACGGCCGCATAATCGAAATTAATACGACTCACTATAGGGGAATTGTGAGCGGATAACAATTCCCC<br>ATCTTAGTATATTAGTTAAGTATAAGAAGGAGATATACATATGAGTGTCAACTTAGCTTCCCAGTT</td></tr>
</table>

| | |
|---|---|
| | GCGGGAAGGGACGAAAAAATCCCACTCCATGGCGGAGAACGTCGGCTTTGTCAAATGCTTCCTC<br>AAGGGCGTTGTCGAGAAAAATTCCTACCGTAAGCTGGTTGGCAATCTCTACTTTGTCTACAGTGC<br>CATGGAAGAGGAAATGGCAAAATTTAAGGACCATCCCATCCTCAGCCACATTTACTTCCCCGAAC<br>TCAACCGCAAACAAAGCCTAGAGCAAGACCTGCAATTCTATTACGGCTCCAACTGGCGGCAAGA<br>AGTGAAAATTTCTGCCGCTGGCCAAGCCTATGTGGACCGAGTCCGGCAAGTGGCCGCTACGGC<br>CCCTGAATTGTTGGTGGCCCATTCCTACACCCGTTACCTGGGGGATCTTTCCGGCGGTCAAATTC<br>TCAAGAAAATTGCCCAAAATGCCATGAATCTCCACGATGGTGGCACAGCTTTCTATGAATTTGCC<br>GACATTGATGACGAAAAGGCTTTTAAAAATACCTACCGTCAAGCTATGAATGATCTGCCCATTGAC<br>CAAGCCACCGCCGAACGGATTGTGGATGAAGCCAATGACGCCTTTGCCATGAACATGAAAATGT<br>TCAACGAACTTGAAGGCAACCTGATCAAGGCGATCGGCATTATGGTGTTCAACAGCCTCACCCGT<br>CGCCGCAGTCAAGGCAGCACCGAAGTTGGCCTCGCCACCTCCGAAGGCTAGAAGCTTGGATCC<br>ATTAAAGAGGAGAAACCCGGGCATGGCCGTCACTGATTTAAGTTTGACCAATTCTTCCCTGATGC<br>CTACGTTGAACCCGATGATTCAACAGTTGGCCCTGGCGATCGCCGCTAGTTGGCAAAGTTTACC<br>CCTCAAGCCCTATCAATTGCCGGAGGATTTGGGCTACGTAGAAGGCCGCCTGGAAGGGGAAAAG<br>TTAGTGATTGAAAATCGGTGCTACCAAACGCCCCAGTTTCGCAAAATGCATTTGGAGTTGGCCAA<br>GGTGGGCAAAGGGTTGGATATTCTCCACTGTGTAATGTTTCCTGAGCCTTTATACGGTCTACCTT<br>TGTTTGGCTGTGACATTGTGGCCGGCCCCGGTGGAGTAAGTGCGGCTATTGCGGATCTATCCCC<br>CACCCAAAGCGATCGCCAATTGCCCGCAGCGTACCAAAAATCATTGGCAGAGCTAGGCCAGCCA<br>GAATTTGAGCAACAACGGGAATTGCCCCCCTGGGGAGAAATATTTTCTGAATATTGTTTATTCATC<br>CGTCCCAGCAATGTCACTGAAGAAGAAAGATTTGTACAAAGGGTAGTGGACTTTTTGCAAATTCA<br>TTGTCACCAATCCATCGTTGCCGAACCCTTGTCTGAAGCTCAAACTTTGGAGCACCGTCAGGGGC<br>AAATTCATTACTGCCAACAACAACAGAAAAATGATAAAACCCGTCGGGTACTGGAAAAAGCTTTTG<br>GGGAAGCTTGGGCGGAACGGTATATGAGCCAAGTCTTATTTGATGTTATCCAATAAGACGTCGGT<br>ACCCTCGAGTCTGGTAAAGAAACCGCTGCTGCGAAAT TTGAACGCCA… |
| **moxGFP-PIF6 (pMH1411)**<br><br>T7 promotor<br>moxGFP<br>PIF6<br>His$_6$-Tag<br>Backbone: pRSET (Invitrogen) | …CCCGCGAAATTAATACGACTCACTATAGGGAGACCACAACGGTTTCCCTCTAGAAATAATTTTG<br>TTTAACTTTAAGAAGGAGATATACATATGGTGTCCAAGGGCGAGGAGCTGTTCACCGGGGTGGT<br>GCCCATCCTGGTCGAGCTGGACGGCGACGTAAACGGCCACAAGTTCTCCGTGCGGGGCGAGGG<br>CGAGGGCGATGCCACCAACGGCAAGCTGACCCTGAAGTTCATCAGCACCACCGGCAAGCTGCC<br>CGTGCCCTGGCCCACCCTCGTGACCACCCTGACCTACGGCGTGCAGAGCTTCTCCCGCTACCC<br>CGACCACATGAAGCGCCACGACTTCTTCAAGAGCGCCATGCCCGAAGGCTACGTCCAGGAGCG<br>CACCATCTCCTTCAAGGACGACGGCACCTACAAGACCCGCGCCGAGGTGAAGTTCGAGGGCGA<br>CACCCTGGTGAACCGCATCGAGCTGAAGGGCATCGACTTCAAGGAGGACGGCAACATCCTGGG<br>GCACAAGCTGGAGTACAACTTCAACTCCCACAACGTCTATATCACCGCCGACAAGCAGAAGAAC<br>GGCATCAAGGCCAACTTCAAGATCCGCCACAACGTGGAGGACGGCTCCGTGCAGCTCGCCGAC<br>CACTACCAGCAGAACACCCCCATCGGCGACGGCCCCGTGCTGCTGCCCGACAACCACTACCTG<br>TCCACCCAGTCCAAGCTGTCCAAAGACCCCAACGAGAAGCGCGATCACATGGTCCTTCTGGAAT<br>TCGTGACCGCCGCCGGGATCACTCACGGCATGGACGAGCTGTACAAGGGCTCCGCAGGTTCTG<br>CTGGTATGATGTTCTTACCAACCGATTACTCGAGCAGGTTAAGCGATCAAGAGTATATGGAGCTT<br>GTGTTTGAGAATGGCCAGATTCTTGCAAAGGGCCAAAGATCCAACGTTTCTCTGCATAATCAACG<br>TACCAAATCGATCATGGATTTGTATGAGGCAGAGTATAACGAGGATTTCATGAAGAGTATCATCCA<br>TGGTGGTGGTGGTGCCATCACAAATCTCGGGGACACGCAGGTTGTTCCACAAAGTCATGTTGCT<br>GCTGCCCATGAAACAAACATGTTGGAAAGCAATAAACATGTTGACCATCATCACCATCACCATTAA<br>AAGCTTGATC… |
| **mCherry-PIF6 (pMH613)**<br><br>T7 promotor | …CCCGCGAAATTAATACGACTCACTATAGGGAGACCACAACGGTTTCCCTCTAGAAATAATTTTG<br>TTTAACTTTAAGAAGGAGATATACATATGGGTTGTCGCGACGGTAGCGCCGGCGTGAGCAAGGG<br>CGAGGAGGATAACATGGCCATCATCAAGGAGTTCATGCGCTTCAAGGTGCACATGGAGGGCTCC<br>GTGAACGGCCACGAGTTCGAGATCGAGGGCGAGGGCGAGGGCCGCCCCTACGAGGGCACCCA<br>GACCGCCAAGCTGAAGGTGACCAAGGGTGGCCCCCTGCCCTTCGCCTGGGACATCCTGTCCCC<br>TCAGTTCATGTACGGCTCCAAGGCCTACGTGAAGCACCCCGCCGACATCCCCGACTACTTGAAG |

| Legend |
| --- |
| Cystein |
| mCherry |
| PIF6 |
| $His_6$-Tag |
| Backbone: pRSET (Invitrogen) |

CTGTCCTTCCCCGAGGGCTTCAAGTGGGAGCGCGTGATGAACTTCGAGGACGGCGGCGTGGTG
ACCGTGACCCAGGACTCCTCCCTGCAGGACGGCGAGTTCATCTACAAGGTGAAGCTGCGCGGC
ACCAACTTCCCCTCCGACGGCCCCGTAATGCAGAAGAAGACCATGGGCTGGGAGGCCTCCTCC
GAGCGGATGTACCCCGAGGACGGCGCCCTGAAGGGCGAGATCAAGCAGAGGCTGAAGCTGAA
GGACGGCGGCCACTACGACGCTGAGGTCAAGACCACCTACAAGGCCAAGAAGCCCGTGCAGCT
GCCCGGCGCCTACAACGTCAACATCAAGTTGGACATCACCTCCCACAACGAGGACTACACCATC
GTGGAACAGTACGAACGCGCCGAGGGCCGCCACTCCACCGGCGGCATGGACGAGCTGTACAAG
GGCTCCGCAGGTTCTGCTGGTATGATGTTCTTACCAACCGATTACTCGAGCAGGTTAAGCGATCA
AGAGTATATGGAGCTTGTGTTTGAGAATGGCCAGATTCTTGCAAAGGGCCAAAGATCCAACGTTT
CTCTGCATAATCAACGTACCAAATCGATCATGGATTTGTATGAGGCAGAGTATAACGAGGATTTCA
TGAAGAGTATCATCCATGGTGGTGGTGGTGCCATCACAAATCTCGGGGACACGCAGGTTGTTCC
ACAAAGTCATGTTGCTGCTGCCCATGAAACAAACATGTTGGAAAGCAATAAACATGTTGACCATC
ATCACCATCACCATTAAAAGCTTGATC…
